# Comprehensive analysis of small RNA profile by massive parallel sequencing in HTLV-1 asymptomatic subjects with monoclonal and polyclonal rearrangement of the T-cell antigen receptor γ-chain

**DOI:** 10.1101/277871

**Authors:** Daniela Raguer Valadão de Souza, Rodrigo Pessôa, Youko Nukui, Juliana Pereira, Jorge Casseb, Augusto César Penalva de Oliveira, Patricia Bianca Clissa, Sabri Saeed Sanabani

## Abstract

**Introduction:** In this study, we used a massive parallel sequencing technology to investigate the cellular small RNA (sRNA) operating in peripheral blood mononuclear cells (PBMCs) of the Human T-lymphotropic virus type I (HTLV-I) infected asymptomatic subjects with a monoclonal and polyclonal rearrangement of the T-cell antigen receptor γ-chain.

**Materials and Methods:** Blood samples from 15 HTLV-1 asymptomatic carriers who were tested for clonal *TCR*-γ gene (seven and eight subjects presented monoclonal and polyclonal expansion of HTLV-1 infected cells, respectively), and were submitted to Illumina for small RNA library construction. sRNA libraries were prepared from cryoperserved PBMCs using TrueSeq Small RNA Library Preparation Kit (Illumina). The sRNA-Seq reads were aligned, annotated, and profiled by different bioinformatics tools.

**Results:** Through bioinformatics analysis, we identified a total of 494 known sRNAs and 120 putative novel sRNAs. Twenty-two known and 15 novel sRNA showed a different expression (>2-fold) between the asymptomatic monoclonal (ASM) and asymptomatic polyclonal carriers (ASP). The hsa-mir-196a-5p was the most abundantly upregulated micro RNA (miRNA) and the hsa-mir-133a followed by hsa-mir-509-3p were significantly downregulated miRNAs with more than a three-fold difference in the ASM than ASP group. The target genes predicted to be regulated by the differentially expressed miRNAs play essential roles in diverse biological processes including cell proliferation, differentiation, and/or apoptosis.

**Discussion:** Our results provide an opportunity for a further understanding of sRNA regulation and function in HTLV-1 infected subjects with monoclonality evidence.

## Introduction

Human T-lymphotropic virus type I (HTLV-I) is an oncogenic human retrovirus that was first isolated from a T-cell line, HUT102, that had been obtained from a patient with adult T-cell leukemia/lymphoma (ATLL) [1]. Globally, there are an estimated 5-10 million individuals who carry the HTLV-I, with the important caveat that the prevalence remains barely unknown in many areas of the world [2]. The disease burden is unevenly distributed, with a higher incidence of the disease particularly in southwest Japan, the Caribbean islands, South America, and portions of Central Africa [3]. Infection with HTLV-I may result in a spectrum of clinical manifestations, ranging from asymptomatic infection to a number of human disorders, most notably a malignant ATLL and a chronic progressive neuromyelopathy, termed HTLV-I-associated myelopathy/tropical spastic paraparesis (HAM/TSP) [4]. The majority of HTLV-I-infected individuals remain asymptomatic for life, while some individuals progress to a pre-leukemic phase that is characterized by small numbers of circulating leukemic cells in the peripheral blood, skin lesions, and a lack of involvement of other organ systems [5]. Only 2.5% to 5% of the virus carriers eventually develop ATLL after a long asymptomatic period [6, 7]. The reason why some people develop the disease, whereas others remain healthy, is likely dependent on both host-related and virus-related factors [8]. Available evidence from molecular studies indicates that the impairment of various cellular functions by viral genes (e.g., *tax* and *HBZ*), genetic and epigenetic changes including DNA methylation, and the host immune system may contribute to the leukemogenesis of ATLL [9-11]. Despite the effective immortalization of the T cells, the markedly prolonged incubation period (>30 years) prior to the onset of ATLL, suggests an additional acquisition of genetic changes beside the viral infection to contribute to the pathogenesis [12].

Infections with HTLV-1 are expected to produce an initial polyclonal T-cell proliferation followed by a monoclonal malignant transformation in asymptomatic HTLV-1 carriers before the diagnosis of Adult T-cell Leukaemia/Lymphoma becomes evident. HTLV-1carriers who have monoclonal integration of HTLV-1 provirus DNA in their mononuclear cells are suggested to be a high-risk group for development of ATLL, but their prognosis varies from being stable long-term carriers to experiencing the development of ATLL [13-16]. Carvalho and Da Fonseca Porto [17] also found a correlation between monoclonal integration of provirus DNA and abnormal lymphocytes in peripheral blood, with a trend for greater severity of the parasitic infection.

Over the past several years, some studies have provided significant evidence to support transcription of many non-protein-coding regions of the mammalian genome [18], yielding a complex network of transcripts that include tremendous numbers of non-coding RNAs (ncRNA). These molecules play significant roles in normal biological processes and in a variety of human diseases [19]. Within this diverse menagerie of ncRNA, small RNAs (sRNAs), have emerged as potential posttranscriptional regulators of gene expression in both eukaryotes and prokaryotes [20]. Because these molecules are highly complex in terms of structural diversity and function, their families have been grouped into structural and regulatory ncRNA [21]. The structural ncRNA includes transfer RNA (tRNA) and ribosomal RNA (rRNA), as well as other small but stable noncoding RNAs, such as small nuclear RNAs (snRNAs), small nucleolar RNAs (snoRNAs), small cytoplasmic RNA (scRNA), Ribonuclease P (RNase P), mitochondrial RNA processing (MRP) RNA, signal recognition particle (SRP) RNA, and telomerase RNA. Regulatory ncRNAs include microRNAs (miRNAs), PIWI-interacting RNAs (piRNAs), and long ncRNAs (lncRNAs) [22]. The miRNAs are perhaps the single best-understood subclass. They are typically 18-25 nucleotides (nt) in length, single-stranded RNAs that have emerged as principal post-transcriptional regulators of gene expression and play a vital role in several cell processes including cell proliferation, differentiation, apoptosis, embryonic development, and tissue differentiation [23, 24]. The mature molecules of miRNAs are assembled into ribonucleoprotein complexes called miRNA-induced silencing complexes (RISC), which inhibits gene expression by perfect complementary binding, for mRNA degradation, or imperfect binding at the 3’UTR region, to inhibit translation[25]. The rigid control of miRNA expression is critically important to maintain cells in normal physiological states [26], while overexpression of miRNAs has been associated with the occurrence and development of various diseases [25, 27]. It has been proven that oncogenic (onco-miRs) and tumor-suppressor (TS-miRs) miRNAs exert control over important cell cycle components and thus results in either acceleration or deceleration of the cell cycle [28-30].

In theory, dysregulation of host cells miRNAs by HTLV-1 might influence the development of ATLL. Indeed, some reports attest to the critical role of cellular miRNA to proliferation and survival of HTLV-1-infected T-cells. Hybridization-based methodologies, such as microarray and PCR-based assays have been used thus far to identify and profile the cellular miRNAs in HTLV-1 infected cell lines [31, 32]. The study by Pichler *et al.* [33], who first used quantitative PCR to study the interconnections between HTLV-1 and cellular miRNAs, confirmed that HTLV-1 transforms host cells by inducing dysregulation of the expression of specific miRNAs, including miR-146a, which is upregulated by Tax protein (an oncoprotein of HTLV-1). Subsequently, other expression studies revealed both up- and downregulation in a number of miRNAs in HTLV-1/ATLL cell lines and primary ATLL cells [34, 35]. Recently, Ruggero and colleagues [36] employed massive sequencing to identify the repertoire of microRNAs and tRNA fragments (tRFs) expressed in HTLV-1-infected cells compared to normal CD4+ T cells. Most of the previous studies mainly focused on expression of specific or diverse miRNA in HTLV-1 infected cell lines, while few studies have focused on determining the expression profiles of sRNAs isolated from human blood samples. Thus, it would be clinically relevant to investigate early gene regulatory mechanisms associated with cell transformation in HTLV-1 asymptomatic patients with monoclonal expansion features.

In this study, we employed Illumina massive parallel sequencing technology to comprehensively characterize the sRNA expression profiles in the peripheral blood mononuclear cells (PBMCs) of HTLV-1 infected asymptomatic subjects with monoclonal (ASM) and polyclonal (ASP) T cell expansions. Our results revealed 494 known sRNAs and 120 putative novel sRNAs. Moreover, 23 known and 15 novel sRNA showed different expressions (>2-fold) between the ASM and the ASP groups. The data obtained in this study provides considerable insight into understanding the expression characteristic of sRNAs in HTLV-1 carriers with monoclonal T cell population. Differentially expressed sRNAs described here may be associated with the transformation of T cells.

## Materials and Methods

### Patients’ characteristics and sample preparations

Peripheral blood samples were initially collected from a relatively large group of asymptomatic HTLV-1 carriers (n = 367). Of these subjects, 15 carriers were selected for this study on the basis of clonality results and classified to ASM (n= 7) and ASP (n=8). The samples were collected after approval by the institutional health research ethic authority and the signing of written informed was obtained from all individual participants included in the study. Isolation of PBMCs was carried out by Ficoll-Hypaque (Amersham, Upsala, Sweden) density centrifugation, washed twice in RPMI 1640 with 10 % fetal calf serum, and stored in liquid nitrogen until use. A summary of the clinical characteristics of study subjects is shown in **Table 1**.

**Table 1.** Demographic and clinical characteristics of HTLV-1 infected asymptomatic carriers subjected to small RNA analysis.

| Sample | Sex | Age | Clonality features | Proviral load (copies/1000 PBMCs) | Total input reads | Total filtered reads (%) |
| --- | --- | --- | --- | --- | --- | --- |
| 131 | Female | 52 | Polyclonal | 17 | 3,944,199 | 165,061 (4,18%) |
| 146 | Female | 59 | Polyclonal | 208 | 12,538,011 | 502,457 (4,01%) |
| 151 | Female | 72 | Polyclonal | 1 | 11,568,287 | 389,210 (3,36%) |
| 152 | Female | 34 | Polyclonal | 60 | 6,323,875 | 479,958 (7,59%) |
| 167 | Female | 58 | Polyclonal | 32 | 5,828,992 | 469,333 (8,05%) |
| 172 | Female | 31 | Polyclonal | 102 | 4,307,133 | 314,845 (7,31%) |
| 182 | Female | 80 | Polyclonal | 22 | 7,195,318 | 350,231 (4,87%) |
| 188 | Female | 49 | Polyclonal | 8 | 8,913,900 | 314,436 (3,53%) |
| 54 | Female | 42 | Monoclonal | ND | 6,452,823 | 54,089 (0,84%) |
| 75 | Female | 45 | Monoclonal | ND | 11,154,254 | 70,707 (0,63%) |
| 124 | Male | 70 | Monoclonal | 327 | 8,113,493 | 142,191 (1,75%) |
| 143 | Female | 65 | Monoclonal | 71 | 5,719,645 | 246,809 (4,32%) |
| 154 | Female | 28 | Monoclonal | 163 | 11,018,111 | 425,570 (3,86%) |
| 200 | Female | 74 | Monoclonal | ND | 7,918,316 | 409,349 (5,17%) |
| 212 | Male | 61 | Monoclonal | 27 | 12,777,420 | 417,351 (3,27%) |

### Genomic DNA and RNA extraction

Genomic DNA was extracted from PBMCs using the QIAamp blood kit (QIAGEN, Tokyo, Japan). The miRNeasy Mini Kit (Qiagen, Hilden, Germany) in conjunction with the TRIzol (Life Technologies, USA) procedure were used to extract total RNA and sRNA following the manufacturer’s protocols. Briefly, 700 μL of TRIzol was added to 200 μL of cryoperserved PBMCs followed by incubation at room temperature for 5 min. Chloroform (Sigma-Aldrich, St. Louis, MO, USA) was added, and the samples were vortexed and incubated at room temperature for 5 min followed by centrifugation at 12,000×*g* for 15 min at 4° C. The aqueous phase containing RNA was transferred to a new tube and isopropanol (Fisher-Scientific, Thermo Fisher Scientific, Waltham, MA, USA) was added. The samples were incubated at room temperature for 10 min followed by centrifugation at 12,000x*g* for 15 min at 4° C. The pellet was washed with 75% Ethanol (Sigma-Aldrich), air-dried, and resuspended in 20 μL nuclease-free water (Ambion). For addition of carriers, 2 μL of glycogen (5 μg/μL) was added during the isopropanol precipitation step. For the miRNeasy Mini Kit, the aqueous phase was added to 1.5 volumes of 100% ethanol, and the mixture was mixed thoroughly by pipetting. The supernatants were then transferred to the miRNeasy Mini spin column by centrifugation. Thereafter, the column was washed with 700 μl of RWT, 500 μl of RPE, and 500 μl of 80% ethanol, and then dried by centrifugation at 12 000×*g* for 5 min. Finally, the sRNAs on the membrane were eluted in 22 μl of RNase-free water and stored at -80° C until further use.

### Concentration measurement of genetic materials

Both DNA and RNA including sRNA concentration measurements by fluorimetry were done on Qubit 2.0 fluorometer using Qubit^®^ DNA or RNA HS Assay Kit (Thermo Fisher Scientific), respectively. Briefly, 10 μL standards or diluted genetic materials were mixed with Qubit^®^ Working Solution and incubated for 2 min. Two standards were prepared, one has 0 ng/μL and the other has 10 ng/μL concentrations. To obtain the calibration curve, the relative fluorescence was plotted against the concentrations of the two standards. The fluorometer built-in software was used to compute the concentration values of the genetic materials of each sample.

### HTLV-1 proviral load determination

The extracted DNA was used as a template to amplify a 97-bp fragment from the HTLV-1 *tax* region using previously published primers [37] and protocol [38]. Amplification and analysis were performed with the Applied Biosystems 7500 real-time PCR system. The standard curves for HTLV-1 *tax* were generated from MT-2 cells of log_10_ dilutions (from 10^5^ to 10^0^ copies). The threshold cycle for each clinical sample was calculated by defining the point at which the fluorescence exceeded a threshold limit. Each sample was assayed in duplicate, and the mean of the two values was considered the copy number of the sample. The HTLV-1 proviral load was calculated as the copy number of HTLV-1 (*tax*) per 1000 cells◻=◻(copy number of HTLV-1 *tax*)/(copy number of *RNase P* gene/2)×1000 cells. The method could detect one copy per 10^3^ PBMCs.

### Analysis of T cell receptor yTCR genes

A DNA-based polymerase chain reaction (PCR) of rearranged yTCR genes was performed according to the previously described protocol [39]. All patients’ PCR products were analyzed with the 3130 ABI Prism capillary electrophoresis equipment. 0.5 μl ROX, 13 μl Hidi, and 1 μl template DNA sample were added to each well in a 96-well plate. Data were analyzed using Genescan and Genotyper software (Applied Biosystem, Foster City, CA). T cell clonalities were blindly determined by visual examination of the electropherograms by two analysts and further confirmed by an expert hematology pathologist (coauthor JP).

### sRNA construction and massive parallel sequencing (MPS)

For each sample in both groups, sRNA libraries were prepared with the Small RNA v1.5 sample preparation kit as per the manufacturer’s instructions (Illumina, San Diego, CA) and previous protocol [40]. Briefly, 5 μl of purified total RNAs were ligated with 1 μl RNA 3′ Adapter and then with a 5′ RNA adapter (Illumina, San Diego, CA). The 5′ adapter also included the sequencing primer. After RT-PCR amplification, the resulting products were analyzed using polyacrylamide gel electrophoresis (PAGE) (6% Novex Tris-borate-EDTA [TBE] PAGE; Invitrogen). After gel electrophoresis, sRNA bands at sizes 145-150 bp were excised and purified. Finally, each four libraries were pooled and up to 8-10 pM of the pooled libraries were loaded and sequenced on the MiSeq platform (Illumina) with a 36 base single-end protocol, according to the manufacturer’s instructions.

### sRNA Data analysis and interpretation

Base calling, demultiplexing, and trimmed FASTQ files were generated using the MiSeq reporter. Only high quality reads with score >30 on the Sanger scale were considered for further analysis. The reads were aligned against the whole genome build: hg19 using Strand NGS v3.1. The later software package was also used for analysis of novel molecule discoveries and interpretations. The distributions of the sRNA data in each clinical condition were conducted according to the quantile normalization algorithm, with a baseline transformation set to the median of all samples. Novel sRNA were discovered and classified by the decision tree method with three-fold validation accuracy using model previously described by Langenberger *et al.* [41]. Additionally, only sRNA sequences meeting the minimum read coverage criterion of >20 were considered as novel or known sRNA and were included in further analyses. sRNAs with fold change more than 2.0 were considered to be differentially expressed. All the sRNA raw data generated by MPS has been deposited to the Zenodo repository.

### Functional annotation and pathway analysis of miRNA target genes

The target genes from differentially expressed miRNAs were predicted by PicTar, TarBase microRNAorg databases. Pathway analysis was carried out for a functional analysis of mapping genes to KEGG pathways [42]. Prediction was performed by using DIANA-microT-CDS [43]. The biological functions of these target genes were determined by the enriched GO analysis (false discovery rate *p*-value <0.05).

## Results

### Patient Characteristics

The characteristics of the subjects in both groups are shown in **Table 1**. All eight participants within the ASP group were female and the median age was 58 years (range, 31-80 years). Six females and one male were assigned to the ASM group and the median age of the entire group of subjects was 61 years (range, 28-74 years). HTLV-1 proviral load levels varied from one copy to 208 copies/10^3^ PBMCs in the ASP group and varied from undetectable to 327 copies/10^3^ PBMCs in the ASM group (two-tailed *P* value = 0.7).

### Description of whole genome sRNA-sequencing data from both groups

Illumina MPS revealed a total of 130,415,317 reads. After removal of low quality data and reads that failed vendor`s QC, 125,435,316 reads were retained. The filtered sRNA reads on quality metrics for each sample used in the alignment to the human genome sequence dataset are shown in **Table 1**. For instance, sample 131 within the ASP group had 3,779,138 filtered sRNA reads out of 3,944,199 of raw reads and sample 143 in the ASM group displayed 5,472,836 of filtered sRNA reads out of 5,719,645 raw reads. We next investigated the percentage of reads assigned to abundant sRNA categories or sRNA biotypes (**Figure 1**, representative for samples 131 and 143). More than 69% and 59% of the reads in samples 131 and 143, respectively, matched to mature miRNA. The rest of the sequences were matched into other types of sRNAs including intronic RNAs, exonic RNAs, scRNAs, snRNAs, snoRNAs, tRNAs, and others. Next, we evaluated the location of these tags on the chromosome. In the majority of samples from both groups, most of the unique tags were located on chromosome 3, 8, 12, and 19 (**table S1**). The Illumina MPS yielded 614 sRNA molecules, of which 494 sequences were derived from known sRNA, and 120 were novel sRNA. Of the 494 known sRNA, 219 were miRNA, 30 scRNA, 9 scRNA_pseudogene, 162 snoRNA, 23 snRNA, and 51 tRNA (**table S2**).Of the 120 novel genes, 25 were novel miRNA, 50 snoRNA, 3 snRNA, 9 tRNA, and 33 unknown (**table S3**). Considering -3p and -5p mature forms of miRNA, our analysis revealed a total of 2104 mature miRNA, of which 22 had had greater than a two-fold change in expression (**table 2).**

**Table 2.**
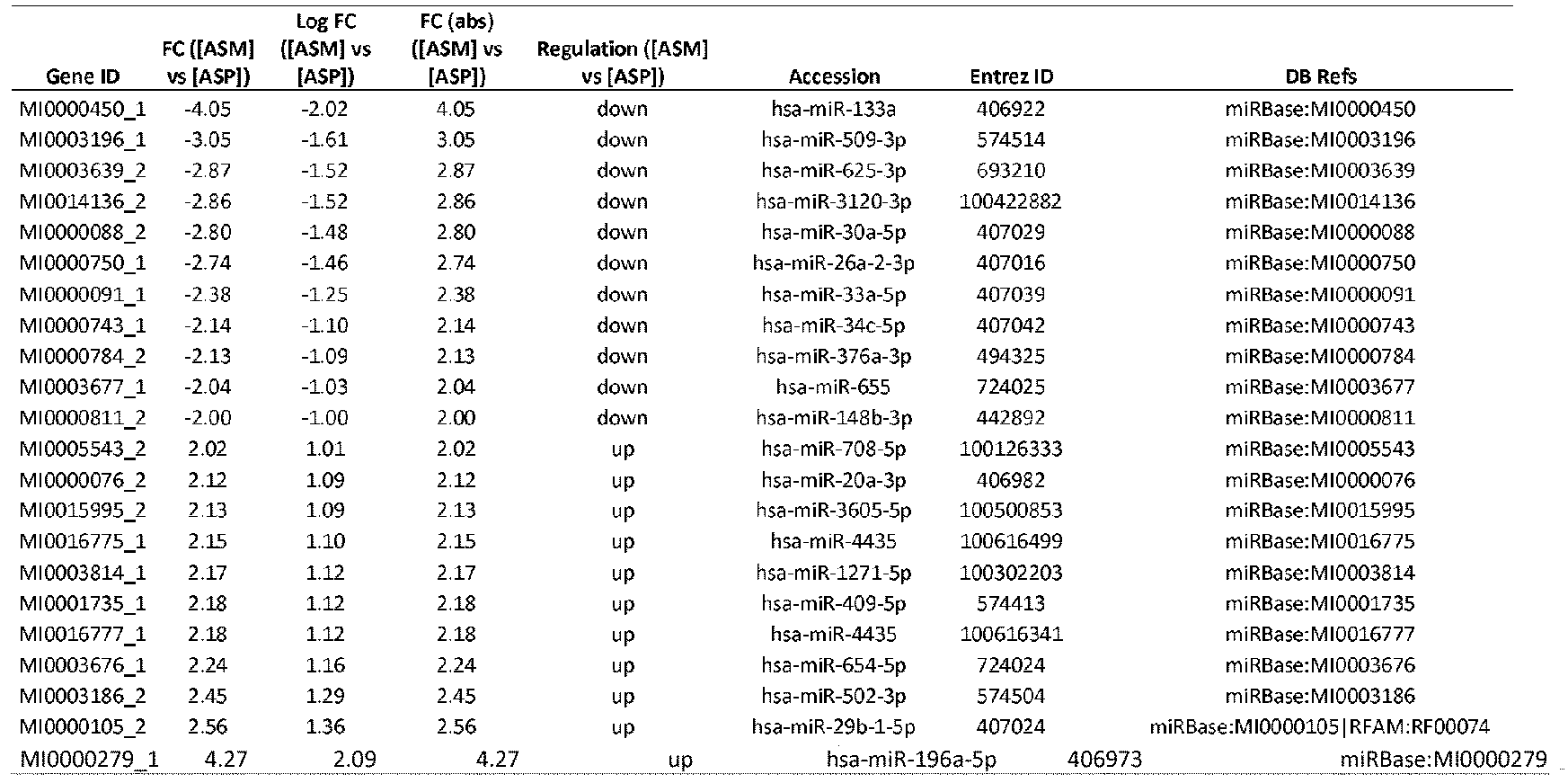
List of miRNAs differentially expressed in asymptomatic HTLV-1 subjects with monoclonal (ASM) vs subjects with polyclonal T cell expansion (ASP)

**Figure 1.**
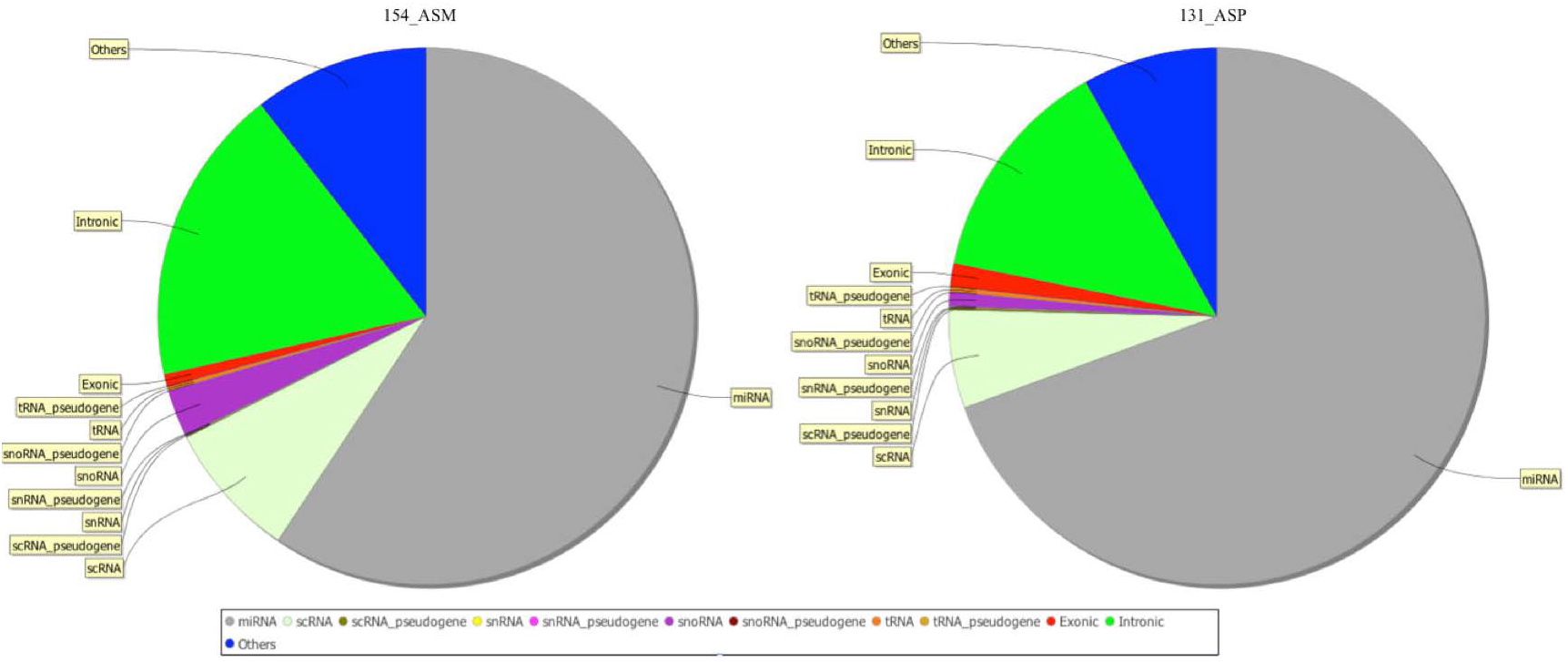
Distribution of the total small RNAs on two representative samples. The clean reads were annotated as tRNAs, rRNAs, snoRNAs, snRNAs, scRNA miRNAs, intro, and others based on the Rfam database

### Characteristics of the abundant known and novel sRNAs

Among the 494 known sRNAs identified in this study, 52 sRNAs were found to be differentially expressed (>2-fold) between the ASM group and ASP group, which accounted for 8.5% (52/614) of all the sRNAs investigated (**table S4**). Clustering by group was not noted in the hierarchal combined tree of the 52 significantly differentially expressed sRNA levels as shown in **figure 2**. Of these 52 sRNAs, 50 were upregulated and two were downregulated in the ASM group. The upregulated sRNA includes three miRNA, 10 scRNA, one scRNA_pseudogene, 21 snoRNA, one snRNA, and 14 tRNA. On the other hand, the down-regulated entities include only two miRNA. Of the highly expressed sRNAs, SNORD45, SNORD45B, hsa-mir-196a-1, hsa-mir-196a-2, and SCARNA5 were the top five most abundantly expressed sRNA in PBMCs of the ASM group, among which, SNORD45 had greater than 5.7 fold changes in expression (**table S4**). The most abundantly downregulated sRNAs in PBMCs of the ASM group were hsa-mir-133a-1 and hsa-mir-133a-2, which showed more than 3.8 fold changes in the ASM when compared to the ASP group (**table S4**). Further, of the 22 mature miRNAs that exhibited a >2-fold difference in expression in the PBMCs of the ASM compared to the ASP group, 11 miRNAs were upregulated and 11 were downregulated (**table 2**). The hierarchical clustering analysis of these miRNAs showed distinct patterns of miRNA expression levels between ASM and ASP groups (**Figure 3**). The hsa-mir-196a-5p was the most abundantly upregulated miRNA and the hsa-mir-133a followed by hsa-mir-509-3p were the significantly downregulated miRNAs with more than a three-fold difference in the ASM than the ASP group. Of the 120 novel sRNA described in this study and shown in **table S3**, 15 were found to be differentially expressed (>2-fold) between the ASM group and ASP group, which accounted for 2.4% of all the sRNAs investigated. Of these, 14 were upregulated and one was downregulated in the PBMCs of the ASM group when compared to the ASP group. Of the upregulated novel sRNAs, four were snoRNA, one miRNA, one tRNA, and eight were unknown while the unique downregulated gene was snoRNA (**Table 3**).

**Table 3.** List of novel small RNAs differentially expressed in asymptomatic HTLV-1 infected subjects with monoclonal (ASM) vs subjects with polyclonal T cell expansion (ASP)

| Gene ID | FC ([ASM] vs [ASP]) | Log FC ([ASM] vs [ASP]) | FC (abs) ([ASM] vs [ASP]) | Regulation ([ASM] vs [ASP]) | Gene Type |
| --- | --- | --- | --- | --- | --- |
| NEWGENE31 | 2.76 | 1.47 | 2.76 | up | Unknown |
| NEWGENE465 | 2.42 | 1.27 | 2.42 | up | snoRNA |
| NEWGENE24 | 2.38 | 1.25 | 2.38 | up | Unknown |
| NEWGENE53 | 2.36 | 1.24 | 2.36 | up | Unknown |
| NEWGENE75 | 2.34 | 1.22 | 2.34 | up | Unknown |
| NEWGENE96 | 2.30 | 1.20 | 2.30 | up | Unknown |
| NEWGENE38 | 2.30 | 1.20 | 2.30 | up | Unknown |
| NEWGENE95 | 2.29 | 1.20 | 2.29 | up | Unknown |
| NEWGENE467 | 2.29 | 1.19 | 2.29 | up | miRNA |
| NEWGENE203 | 2.25 | 1.17 | 2.25 | up | snoRNA |
| NEWGENE377 | 2.23 | 1.16 | 2.23 | up | snoRNA |
| NEWGENE466 | 2.22 | 1.15 | 2.22 | up | tRNA |
| NEWGENE332 | 2.12 | 1.09 | 2.12 | up | snoRNA |
| NEWGENE48 | 2.02 | 1.02 | 2.02 | up | Unknown |
| NEWGENE186 | -2.03 | -0.10 | 2.03 | down | snoRNA |

**Figure 2.**
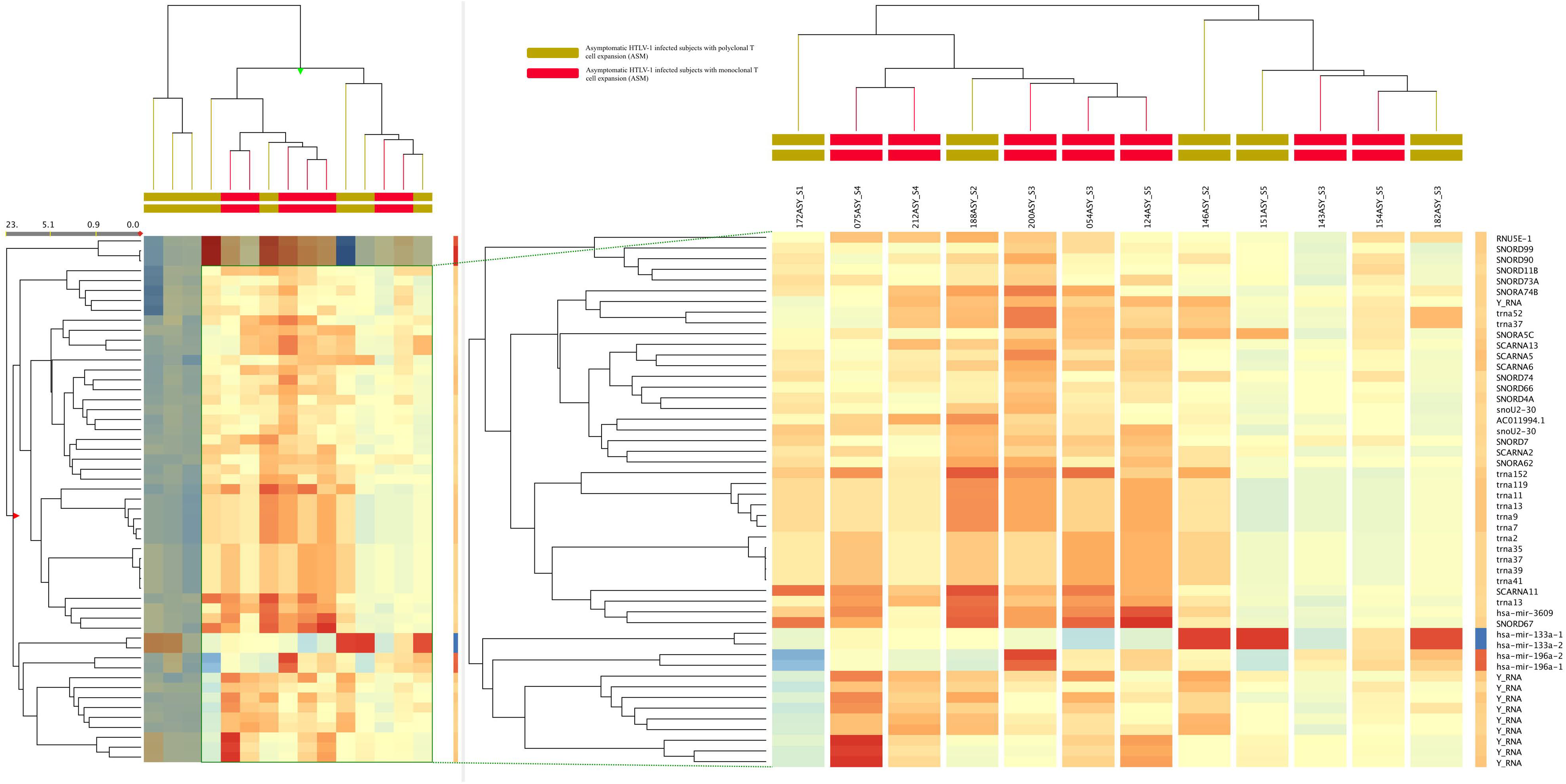
Hierarchical cluster analysis of the 52 known small RNAs differentially expressed comparing ASM vs ASP samples. Rows of the heatmap represent the 52 small RNAs, columns correspond to samples. The cells are colored based on the deviance of the small RNAs expression in the sample from the average expression of the small RNAs, therefore red and blue cells represent respectively expression values higher or lower than mean expression across all samples (white) with color intensity proportional to the difference from the mean, in the regularized logarithmic scale. For clarity purposes, only a subimage of the heatmap has been shown in details.

**Figure 3.**
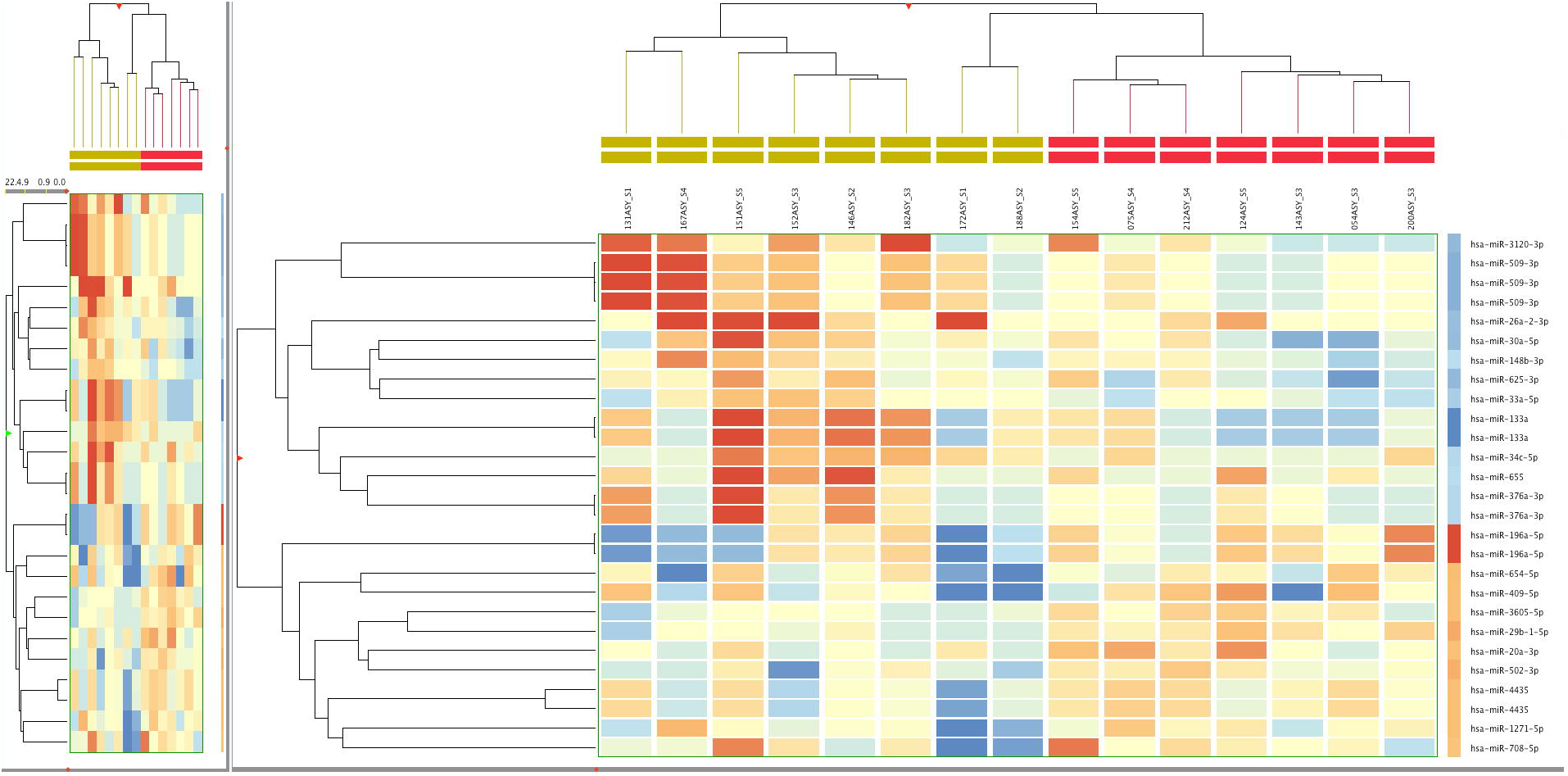
Hierarchical cluster analysis of miRNAs differentially expressed comparing ASM vs ASP samples. Hierarchical clusters were generated using the Euclidean distance and Pearson correlations of miRNA expression patterns. The up-(red) and down-regulated (blue) miRNAs were scaled according to the depicted color code.

### GO Analysis of the Target Genes of the Differentially Expressed sRNAs

The role of a miRNA is ultimately defined by the genes that it targets and by its effect on the expression of those genes. To discover these functions, the target genes of the two groups of the mature significantly expressed miRNAs were predicted using the three programs PicTar, TarBase, and microRNAorg databases. The intersection of these three programs was used to reduce the false-positive rate. As shown in **Table S5**, a *p*-value cutoff of 0.001 was selected to predict nine target genes. The most significantly selected candidate target gene FGD6 is regulated by 13 of the 27 mature significantly expressed miRNAs. The second most selected target gene is the CUGBP Elav-like family member 6 (CELF6) gene, which is regulated by 10 of the 27 mature significantly expressed miRNAs. To identify the actual regulatory functions of the miRNAs, the GO annotation of the target genes was performed on the list of predicted target miRNAs that corresponded with altered miRNAs in the ASM group at p value <0.05. The results from this analysis revealed no significant GO enrichments for the differentially expressed genes predicted from the known miRNAs. The Kyoto Encyclopedia of Genes and Genomes (KEGG) pathways of target genes were assigned using mirPath v.3 in DIANA tools webserver. KEGG pathway annotation revealed that 3044 background genes were annotated for 56 biological functions. The top five pathways were Adherens junction, Proteoglycans in cancer, Axon guidance, Morphine addiction, and Prion diseases.

## Discussion

In this study, we used Illumina high-throughput sequencing technology to analyse the global expression of non-coding RNome in PBMCs of HTLV-1 infected asymptomatic subjects with monoclonal and polyclonal rearrangement of the T-cell antigen receptor γ-chain. Our results revealed 52 significantly differentially expressed sRNAs in the ASM group compared with the ASP subjects. Furthermore, we identified 15 novel sRNA exhibited much higher expression levels in ASM patients than reads in ASP. Eleven of the 22 differentially expressed mature miRNA were up-regulated while the other 11 were down-regulated in the ASM group. Out of the 22 most dysregulated miRs, we selected the most significantly up-regulated miRs (miR-196a-5p) and down-regulated miRs (miR-133a, miR-625-3p, and miR-3120-3p) assuming that the measurement of these miRs could discriminate ASM from ASP patients and that the dysregulation of these miRs in ASM may represent early markers for T cell transformation with unknown functional consequences.

We searched for putative target genes associated with enhancement of cellular transformation. The upregulation of miR-196a, located in HOX gene clusters [44], has been suggested to potentially target HOXB8, HOXC8, HOXD8 and HOXA7 [45]. HOX proteins consist of homeodomain-containing transcription factors that are primary determinants of cell fate during embryogenesis, organogenesis, and oncogenesis [46]. Reports from previous studies have proved that upregulation of miR-196a promotes cell proliferation, anchorage-independent growth, and suppressed apoptosis by targeting annexin A1 [47]. Although most studies proved that over expressed miR-196a is associated with oncogenic function in various types of cancers including pancreatic cancer [48], and oesophageal adenocarcinoma [49] but it has been suggested to play a tumor suppressive role as well. For instance, Li *et al.* [50] have demonstrated a potential role of the up-regulated miR-196a in reducing *in vitro* invasion and *in vivo* spontaneous metastasis of breast cancer cells. Also, miR-196 has been previously demonstrated to play diverse biological functions involving immune response, inflammation, and virus defense [51]. It was also suggested that upregulation of miR-196a may be used in a novel strategy to prevent or treat hepatitis caused by C virus infection, and miR-196a may be valuable in the diagnosis and management of this disease [52, 53]. In relation to T cell leukemia, Coskun and colleagues [54] have shown that miR-196a and miR-196b are regulators of the oncogenic erythroblast transformation-specific (ETS) transcription factors of the ETS related genes (ERG). The ERG plays important physiological and oncogenic roles in hematopoiesis [55]. It is also a prognostic factor in a subset of adult patients with acute T cell leukemia [56]. Coskun and colleagues [54] provided evidence that both miR-196a and miR-196b expression were linked to immature immunophenotype (CD34 positive) in ATLL patients. These findings indicate miR-196a and miR-196b as ERG regulators and have potential implications in ATLL. Based on the above facts, we therefore assumed that the aberrant expression of miR-196a might contribute to the development and transformation of HTLV 1 infected cells in the ASM group.

The miR-133a (known to control HLA-G protein stability) is one of the canonical myomiRs that at the crossroads of the molecular of muscle cells, connecting between pathways for cell differentiation, development and maintenance, but also potentiates aberrant cell growth in a wide range of non-muscle cancers [57]. miR-133 was predicted to bind to both N and G transcripts of rabies viruses and this interaction possibly explains the lengthy dormant state of this virus in skeletal muscles during the early phase of infection [58, 59]. In a recent study, the expression change of miR-133a has been shown to negatively affect the replication of dengue viruses when overexpressed in mammalian cells [60]. The deregulation of miR-133a has been reported in many tumor types of distinct origin, including ovarian [61], prostate [62], bladder [63], and head and neck squamous cell cancer [64]. In most of these cancers, the miR-133a together with other miRs is associated with negative regulation of Fascin. The latter is a global protein that is strongly upregulated in ATLL-derived cells and in CD4+ T-cells transformed by HTLV-1 or by the viral oncoprotein Tax [65]. The loss of miR-133a expression was also associated with a poor survival of tumor patients [66]. The down-regulated miR-133a together with dysregulation of other documented miRs in this study was positively and significantly correlated with some predicted potential target genes including the CUG-binding protein and embryonic lethal abnormal vision-type RNA-binding protein 3-like factor 6 (CELF6), calcium/calmodulin-dependent serine protein kinase interacting protein 2 (CASKIN2), exportin 1 (XPO1), and the solute carrier family 33 member 1 (SLC33A1). The CELF6 protein is an important regulator of cell-specific alternative splicing during normal development and disease [67]. Altered expression of CELF6 has been associated with renal disorders and body weight changes [68, 69]. CASKIN2 is a neuronal signaling scaffolding protein involved in signaling neuronal synapses. In vitro experiments revealed that overexpression of Caskin2 inhibits serum-induced endothelial cells (EC), proliferation, and DNA synthesis but promotes EC survival during serum starvation, thus suggesting a role of this protein in preventing EC dysfunction in vivo [70]. The human chromosomal maintenance 1 or XPO1 mediates the nuclear export of cellular proteins (cargos) bearing a leucine-rich nuclear export signal) and of RNAs. Various viruses, among them HIV-1, HTLV-1 and influenza A use it to export their unspliced or incompletely spliced RNAs out of the nucleus. The XPO1 interacts with, and mediates the nuclear export of HTLV-1 Rex proteins and critically needs it for its multimerization [71, 72]. The SLC33A1 is an important substrate for a large variety of biochemical reactions occurring in the cell [73].

The search for the putative target RNA molecules in the database showed that miR-625-3p can potentially target the gene that codes for the clathrin heavy chain (CLTC) protein. Clathrin is the major protein component of the coat that surrounds the cytoplasmic face of the organelles (coated vesicles) mediating selective protein transport [74]. Clathrin coats play an essential role in receptor-mediated endocytosis, localization of resident membrane proteins to the compartments within the trans-Golgi network, and transport of proteins to the lysosome/vacuole [75, 76]. The clathrin coats contain both clathrin and four adaptor complexes (APs 1-4), which participate in the selection of protein cargo and their incorporation into clathrin-coated transport vesicles [77]. Previous studies showed that AP-2 and AP-3 interact with HIV-1 Gag and contribute to its trafficking and release [78, 79] and that the disruption of this interaction enhances viral release, while reducing the infectivity of the virions produced [78]. Camus and colleagues, 2007 [80] propose that in HIV-1 and murine leukemia virus, the AP-1 promotes Gag release by transporting it to intracellular sites of active budding, and/or by facilitating its interactions with other cellular partners. The same study concluded that this interaction is conserved among different retroviruses including HTLV and RSV and suggested that AP-1 could also play a role in the assembly of these retroviruses. In human disease, the few available data indicate that the abnormalities and dysregulation of CLTC are associated with large B-cell lymphoma [81], inflammatory myofibroblastic tumor [82], and multinodular goiter pathogenesis [83].

miR-3120-3p was down-regulated (fold change = -3.154) in the ASM group, and was negatively correlated with diverse putative target genes such as the SLC33A1, tripartite motif containing 27 (TRIM27), XPO1, CASKIN2, and CELF6. The TRIM27 is a member of a large family of proteins that function as Really Interesting New Gene [82] E3 ubiquitin ligases. The TRIM27 has the ability to inhibit the class II phosphatidylinositol 3 kinase C2β activity, resulting in decreased KCa3.1 channel activity and decreased TCR-stimulated Ca2+ influx and cytokine production, thereby identifying TRIM27 as a unique negative regulator of CD4 T cells [84].

There are few microarrays studies that have also characterized the miRNA expression profiles in HTLV-1/ATLL cell lines and ATLL patients [33-35, 85]; however, it is difficult to compare their results with our study due to the different methods employed, different research designs, and different target population.

Our study has several limitations, particularly its retrospective design, and lack of a control non-infected group. Additionally, few samples were included, and no validation study was performed. In spite of these caveats, the current study unravels a number of known and novel subsets of dysregulated sRNA including signature in the PBMCs of the ASM group. The aberrant expression of sRNAs paves the way for assessment of their mechanistic roles and clinical utilities as biomarkers, prognostic indicators, and/or therapeutic targets in early HTLV-1 pathogenesis.

In conclusion, significantly differentially expressed sRNAs including miRNAs and their target genes between ASM and ASP were identified by means of MPS technique. These newly identified miRNAs and target could have unexplored functions in HTLV-1 pathogenesis. Therefore, further investigation of a large population of leukaemia patients using well-matched control groups and sequential samples will be required to elucidate the possible pathogenic impact of these ncRNA molecules in HTLV-1 infection.

## Acknowledgements

This study was funded by the Fundação de Amparo à Pesquisa do Estado de São Paulo (grants 2011/12297-2 and 2014/24596-2)

## Author Contributions

Conceived and designed the experiments: SSS. Performed the experiments: DRVS, RP, SSS. Analyzed the data: DRVS, RP, PBC, SSS. Contributed reagents/materials/analysis tools: YN JP. Wrote the paper: PBC,SSS. Attending physician for sample collection: YN JP JK ACS.

## Supporting Information

Table S1: Distribution of the small RNA massively sequencuing reads according to the chromosomes

Table S2: A full list of all known small RNAs identified in this study

Table S3: A full list of all novel small RNAs identified in this study

Table S4: List of small RNAs differentially expressed in asymptomatic HTLV-1 infected subjects with monoclonal (ASM) vs subjects with polyclonal T cell expansion (ASP)

Table S5: List of the potential protein targets for the deferentialy expressed miRNA in the ASM group

